# Somatic Genome Diversification and Clonal Evolution of Pathogenic B Cells in Systemic Autoimmunity

**DOI:** 10.64898/2026.06.08.730997

**Authors:** Yumika Shiba, Pathum Kossinna, Loïc Caloren, Lizzy Pijpers, Xuyao Li, Arghavan Ashouri, Jianfan Ivy Nie, Dennisse Bonilla, Laura P. Whittall-Garcia, Joan E. Wither, Dafna D. Gladman, Zahi Touma, Leandro Venturutti, Federico Gaiti

## Abstract

Systemic autoimmune rheumatic diseases are characterized by persistent immune activation and clinical heterogeneity, yet the molecular processes sustaining and diversifying pathogenic immune states remain incompletely understood. Here, we investigate somatic genome diversification as a potential driver of immune variation and disease progression. Using Systemic Lupus Erythematosus as a model disease, we generated a subset-resolved map of somatic mutations across B-cell subsets from 35 patients. Double-negative (DN) B cells carried the highest mutational burden, which was associated with disease duration and immunosuppressive therapy, rather than with disease activity. Integration with single-cell transcriptomes linked DN mutations to dysregulated signalling and proteostasis. Notably, DN cells harboured mutations in genes recurrently altered in B-cell lymphomas. Together, these findings identify somatic genome diversification as a feature of pathogenic B-cell subsets and provide a resource for understanding how chronic stimulation and therapeutic pressure shape the clonal evolution of autoimmunity.

**KEY POINTS:**

- Double-negative B cells in SLE carry the highest somatic mutational burden among circulating B-cell subsets and show evidence of treatment-associated mutational processes.
- Pathogenic variants in DN cells are acquired *de novo* or through selective expansion of pre-existing variant-bearing clones.
- Acquisition of lymphoma-relevant mutations by DN cells is common, supporting a molecular-level linkage between chronic systemic autoimmunity and B-cell malignancies.
- The high-depth, subset-resolved map provides a foundational resource for the functional prioritization of variants and the development of targeted sequencing strategies for monitoring pathogenic clones.

## INTRODUCTION

Systemic autoimmune rheumatic diseases (SARDs) are marked by substantial immunological heterogeneity, leading to highly variable manifestations that complicate diagnosis, risk stratification, and treatment^1^. Disease trajectories are dynamically shaped by the integrated influence of cell-intrinsic molecular programs, immunomodulatory therapies, and cumulative immune injury. Identifying and deconvoluting the contribution of these factors remains a major unresolved challenge. In particular, although inherited genetic variation is a well-established contributor to autoimmune disease susceptibility^2–7^, germline risk alone does not fully account for the marked heterogeneity in disease progression, treatment response, or long-term complications observed across patients. A central unresolved question is therefore whether pathogenic immune cell populations undergo progressive somatic diversification during chronic autoimmune disease, thereby contributing to immune dysfunction, persistence, and clinical heterogeneity.

In hematologic malignancies, somatic evolution drives clonal selection, functional divergence, and disease progression^8^. Emerging evidence suggests that related processes may also shape pathogenic immune populations in non-malignant settings. In mouse models, B cells engineered to harbour canonical lymphoma driver mutations such as *MYD88 L265P* acquire phenotypes resembling human autoimmune cells^9,10^. Conversely, autoreactive B-cell populations in patients with SARDs have been reported to carry mutations in genes recurrently altered in lymphoid malignancies^11^. Recent work in autoimmune thyroid disease further identified convergent acquisition of immune-regulatory driver mutations across independent B-cell clones, supporting the idea that somatic evolution can contribute directly to immune persistence and tolerance escape^12^. More broadly, studies in normal human tissues have shown that somatic genomes can retain durable records of exogenous mutagen exposure, raising the possibility that circulating immune cells can similarly record the cumulative effects of chronic inflammation and therapeutic pressure^13,14^. Together, these observations suggest that somatic mutagenesis may contribute to the emergence, persistence, and/or amplification of pathogenic immune states in chronic systemic autoimmunity^15,16^.

Systemic lupus erythematosus (SLE) is a prototypical B-cell-mediated SARD, characterized by chronic inflammation, immune dysregulation, and multi-organ involvement, making it an ideal model in which to investigate somatic genome diversification in pathogenic immune populations. Clinical manifestations range from mild cutaneous involvement to life-threatening organ dysfunction, highlighting the substantial heterogeneity that defines SLE^17,18^. Patients with SLE are also at increased risk of developing B-cell lymphomas compared to the general population^19^, including aggressive mature B-cell lymphomas such as diffuse large B-cell lymphoma^20^, further suggesting a link between chronic immune dysregulation and aberrant clonal evolution. A consistent immunological feature of SLE is the expansion of antigen-experienced, self-reactive B cells lacking IgD and CD27 expression, termed double-negative (DN) B cells^9,21,22^. DN B cells give rise to autoantibody-secreting plasma cells, and their accumulation correlates with increased SLE disease activity^9,23^. In preclinical models, ablation of DN B cells blocks SLE progression, highlighting their pathogenic relevance^24^. Together, these features position DN B cells as a key pathogenic population in which to assess whether and how somatic genome diversification occurs^8–11,15,16^.

Here, we generated a subset-resolved map of somatic mutations across purified circulating B-cell populations in SLE, including DN, naïve, and memory B cells. Using high-coverage whole-exome sequencing and integrating these data with single-cell transcriptomic profiling, we define the somatic mutational landscape of circulating B-cell subsets and identify DN B cells as a pathogenic compartment with pronounced somatic genome diversification. Our findings provide a framework for understanding how disease-associated B-cell populations emerge and persist in SARDs and for investigating how somatic genome diversification contributes to disease heterogeneity in these autoimmune disorders.

## RESULTS

### Circulating SLE DN cells exhibit increased somatic mutational burden

To define somatic mutational landscapes across circulating B-cell subsets in SLE, we isolated naïve B cells (NB, CD19^+^IgD^+^), memory B cells (MBC, CD19^+^IgD^−^CD27^+^), and double-negative B cells (DN, CD19^+^IgD^−^CD27^−^) from peripheral blood mononuclear cells (PBMCs) of 35 adult SLE patients by fluorescence-activated cell sorting (FACS; **Fig. 1A**). This all-comers cohort captured the expected clinical heterogeneity of SLE, spanning a broad range of disease activity, disease duration, and treatment exposure (**Fig. 1B)**. Disease activity at collection, measured by the SLE Disease Activity Index 2000 Glucocorticoids [SLEDAI-2KG] score^25^ (higher score indicates more severe symptoms), ranged from 0 to 22, with a median [Q1,Q3] of 8 [4,13], whereas time since diagnosis ranged from 3 to 46 years, with a median of 12 years [8,20]. As expected, most patients had received antimalarials, glucocorticoids, and immunosuppressive therapies^26,27^, with smaller subsets also exposed to biologics and/or anticoagulants (**Fig. 1B**).

**Figure 1.**
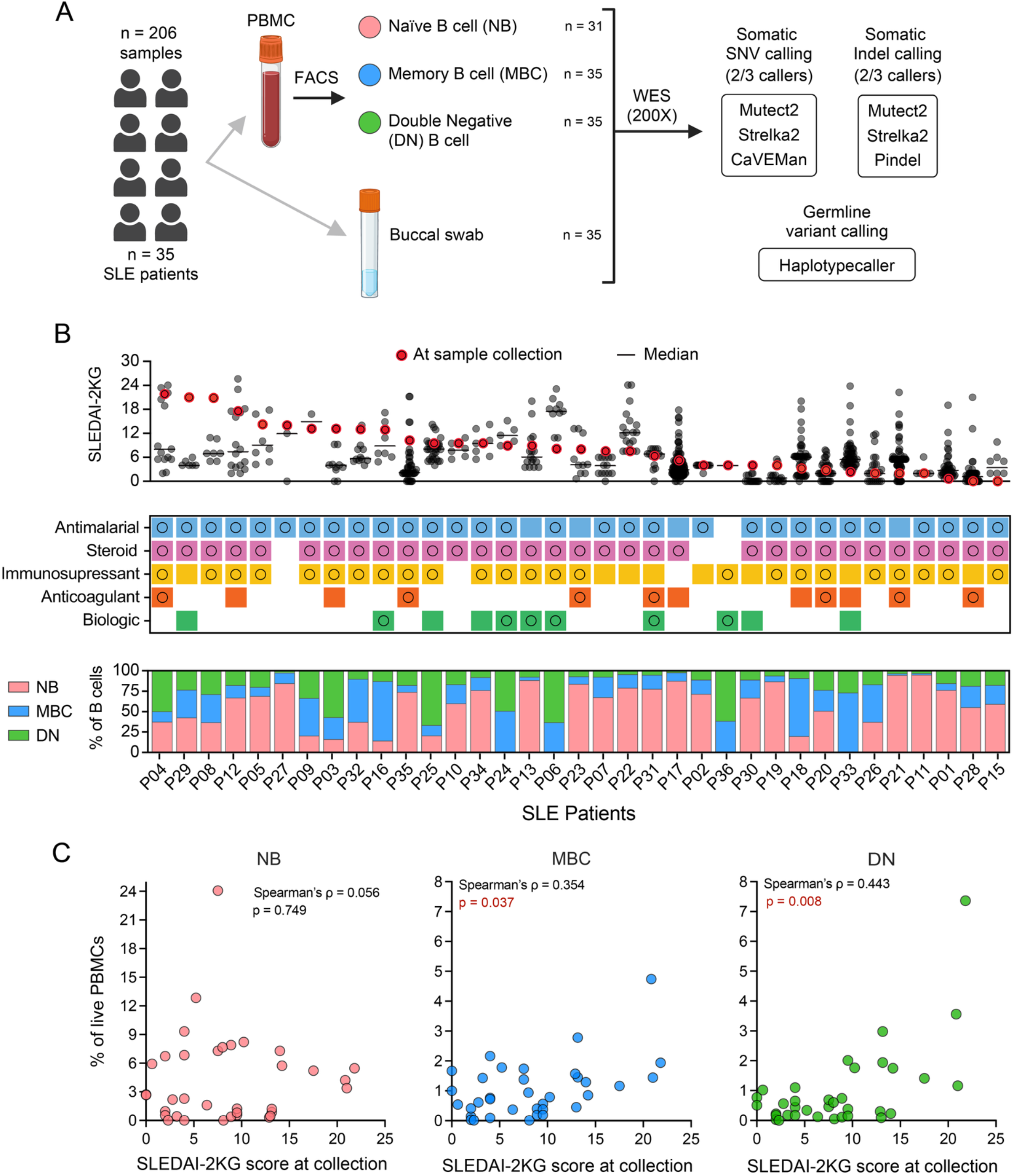
Overview of the SLE patient cohort. **(A)** Study workflow for profiling somatic mutations, including single-nucleotide variants (SNVs) and insertions/deletions (indels), in circulating naïve B cells (NB), memory B cells (MBC), and double-negative B cells (DN) from patients with SLE (n=35). **(B)** Patient-level distribution of clinical treatment and immunophenotypic features. ***Top***, SLE disease activity at collection, measured by SLEDAI-2KG. ***Middle***, treatment exposure. Colours indicate prior exposure to the indicated drug class; circles indicate treatment at the time of sample collection. ***Bottom***, proportions of B-cell subsets among total B cells. Patients are ordered by decreasing SLEDAI-2KG score at collection. **(C)** Proportion of B-cell subsets among live peripheral blood mononuclear cells (PBMCs) in relation to SLEDAI-2KG score at collection. Each point represents one individual patient. Associations were assessed using Spearman rank correlation; Spearman’s rho (ρ) and corresponding *P* values are shown.

In line with prior reports^9^, the proportions of MBC and DN cells among live PBMCs increased with disease activity (ρ=0.354, p=0.037; ρ=0.443, p=0.008, respectively), whereas the abundance of NB did not (**Fig. 1C**). By contrast, none of the three subsets showed a significant association with age, supporting the idea that expansion of DN cells reflects disease-associated immune dysregulation rather than chronological aging^23^.

To identify B-cell-specific somatic variants, we next performed high-coverage whole-exome sequencing (∼200x) across 206 samples, from the 35 patients, including purified B-cell populations, matched non-B-lineage hematopoietic controls, and buccal DNA for germline exclusion (**Fig. 1A**). Somatic single-nucleotide variants (SNVs) and insertions/deletions (indels) were called using a consensus approach whereby only high-confidence variants, supported by at least two variant callers, were retained for downstream analysis (**Methods**).

We observed marked inter-patient variation in somatic mutational burden (**Fig. 2A**). Across all B-cell samples, the median [Q1,Q3] number of SNVs and indels per patient was 98 [23,396] and 19 [7,111], respectively. Deletions accounted for most indels (mean 92.7% per patient; **Fig. 2A**), whereas transitions were the most common SNV class (mean 55.1% per patient; **Fig. 2A**). When stratified by B-cell subset, MBC and DN samples showed significantly higher normalized somatic mutational burden than NB samples, with the highest burden observed in DN cells (**Fig. 2B**). Variant allele frequencies (VAFs) were generally low across all three B-cell compartments, indicating that most somatic variants were present in subclonal fractions of circulating B-cell populations. Together, these results indicate that somatic mutation accumulation is not evenly distributed across circulating B-cell subsets and is concentrated in antigen-experienced populations, particularly DN cells.

**Figure 2.**
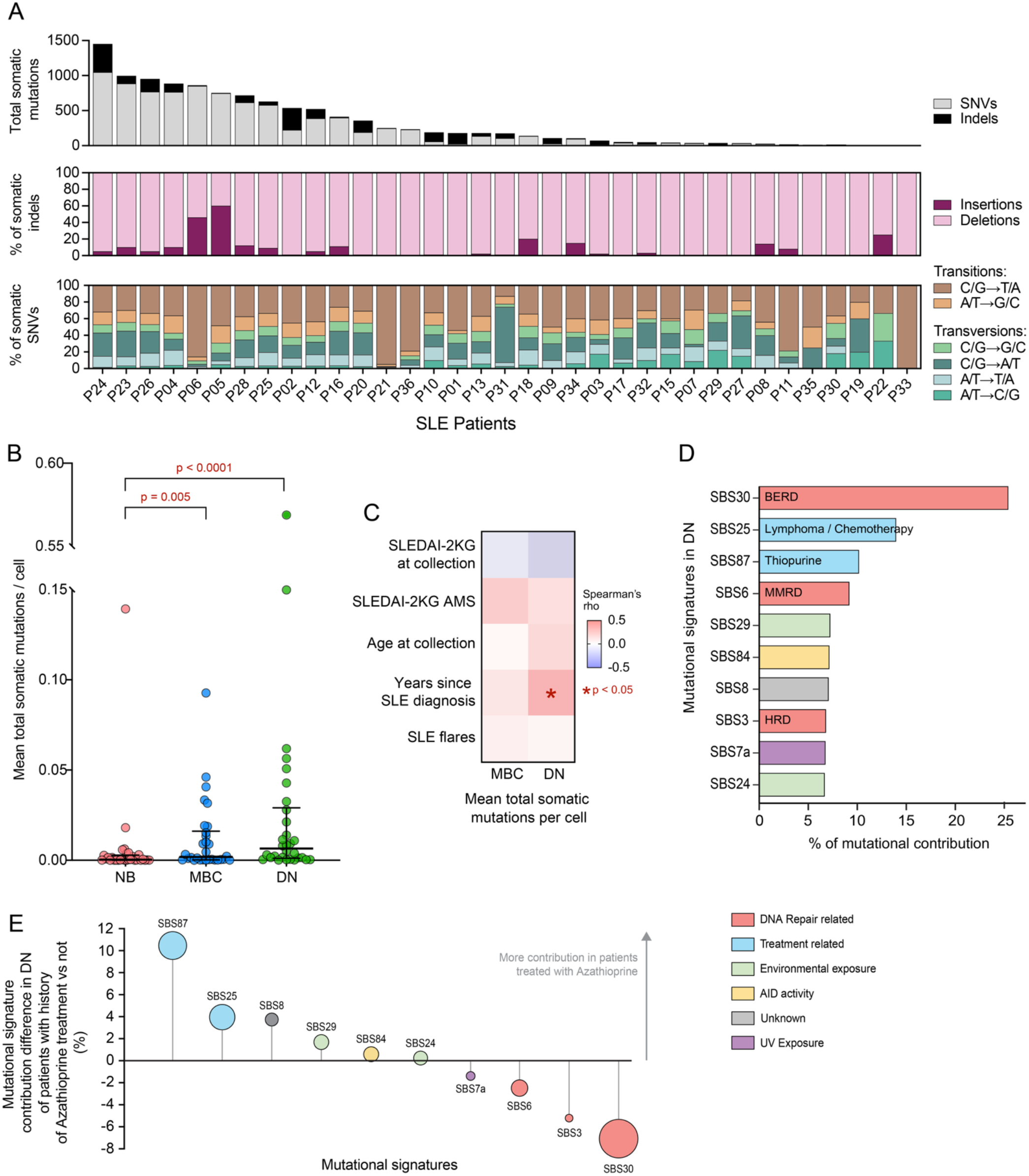
Double-negative (DN) B cells in SLE accumulate more somatic mutations than other B-cell subsets. **(A)** Patient-level distribution of somatic mutational profiles. ***Top***, number of somatic mutations, annotated by mutation type (SNVs or indels). ***Middle***, proportion of insertions and deletions among all somatic indels. ***Bottom***, distribution of transition and transversion classes among all SNVs. Patients are ordered by the total number of somatic mutations (SNVs and indels) detected. **(B)** Cell-number-normalized somatic mutational burden across B-cell subsets. Black bars indicate the median, first quartile (Q1) and third quartile (Q3). For each patient and subset, mutational burden was calculated as the total number of somatic mutations divided by the number of sorted cells in that subset. *P* values were determined by pairwise Dunn’s tests (n=30 patients with matched evaluable NB, MBC, and DN samples). **(C)** Heatmap showing associations between cell-number-normalized somatic mutational burden and the indicated clinical variables. The colour scale represents Spearman’s rho correlation coefficients; statistically significant associations (*P* < 0.05) are marked with an asterisk. Flares were defined as in Touma *et al*.^28^ **(D)** Relative contribution of the top 10 individual mutational signatures in DN cells. Signatures are ranked from highest to lowest contribution. BERD: Base Excision Repair Deficiency; MMRD: Mismatch Repair Deficiency; HRD: Homologous Recombination Deficiency; AID activity: Activation-Induced cytidine Deaminase activity. Signature analysis was restricted to SNVs, as indel counts were too low for robust inference. **(E)** Difference in mean mutational signature contribution in DN cells between patients with and without prior azathioprine exposure. Positive values indicate greater signature contribution in azathioprine-exposed patients. Circle size denotes the mean contribution of each signature across all patients evaluated for DN mutational signatures.

To shed light on the aetiology of the elevated somatic mutational burden in DN cells and, to a lesser extent, in MBC, we next assessed associations with clinical features relevant to SLE. Somatic mutational burden showed no significant association with age, disease activity at collection, cumulative disease activity over time (Adjusted Mean SLEDAI-2KG [AMS]), or number of documented flares^28^ (**Fig. 2C)**. However, disease duration (years since diagnosis) was positively associated with DN mutational burden (ρ=0.357, p=0.035), suggesting that mutation accrual in this compartment reflects longitudinal disease-associated processes and cumulative exposures, rather than acute inflammatory activity alone (**Fig. 2C)**.

To identify processes contributing to these genetic alterations, we further derived mutational signatures. As expected for antigen-experienced B cells, we detected SBS84 in DN and MBC (**Fig. 2D**), consistent with activation-induced cytidine deaminase (AID) activity linked to somatic hypermutation. The clock-like signature SBS1 was also present in both DN and MBC populations, but its relatively modest contribution suggested that age-related mutagenesis was not the primary driver of the increased burden observed in DN cells.

In contrast, DN cells showed greater contribution from signatures linked to DNA damage and repair-related processes (**Fig. 2D**), consistent with the genotoxic milieu associated with SLE^29^. Notably, SBS30, a signature associated with oxidative DNA damage and base-excision repair, accounted for more than 25% of DN mutations and contributed nearly threefold more in DN cells than in MBC, suggesting disproportionate oxidative genotoxic stress in DN cells^30^ (**Fig. 2D**).

Therapy-associated signatures, including SBS87 (thiopurine exposure) and SBS25, were also detected in both DN and MBC but contributed more prominently to the DN landscape (**Fig. 2D**), consistent with prior work showing that treatment exposure can leave durable mutational footprints in non-malignant tissues^31^. In line with this interpretation, among patients with evaluable DN mutational signatures, those with documented prior/ongoing exposure to the thiopurine immunosuppressant azathioprine (Imuran; n=9) showed increased contribution of treatment-associated signatures, particularly SBS87, compared with azathioprine-naïve patients in our cohort (n=6; **Fig. 2E**). These patients also showed enrichment of SBS8, a damage-associated signature whose genomic distribution has recently been shown to vary with DNA repair proficiency and replication stress^32^, suggesting that thiopurine exposure may occur within a broader genotoxic/repair-stress landscape in DN cells.

Together, these findings identify DN B cells as the circulating B-cell subset with the highest somatic mutational burden in SLE and suggest that their mutational landscape integrates cumulative disease-associated genotoxic stress and treatment-related mutagenic exposures.

### DN cells accumulate both subset-specific and selectively expanded shared variants

Given the elevated mutational burden in DN cells and their central pathogenic relevance in SLE, we next sought to define how somatic variants arise and are maintained within this compartment. We distinguished two major classes of alterations: “unique to DN” mutations, which likely reflect genetic alterations acquired after emergence of the DN state, and “shared by DN” mutations, which were presumably acquired by a parental NB/MBC and later inherited by DN.

DN cells harboured the largest number of subset-specific SNVs (**Fig. 3A**), consistent with continued mutation acquisition after differentiation into the DN compartment. Non-synonymous SNVs uniquely detected in DN cells were primarily enriched in pathways related to cell signalling, cell-cycle regulation, and DNA damage responses (**Fig. 3B**), all relevant to DN-cell activation and expansion. Several altered genes within these pathways (e.g., *MTOR, TLR* family genes, and *STAT1*) have documented roles in SLE pathogenesis and immune dysregulation^33,34^, suggesting that additional mutated genes identified here may be similarly relevant to DN-cell pathobiology. Mutations in DNA damage-response genes implicated in B-cell biology, including *PRKDC*^35^ and *XRCC5*^36^, are of particular interest because these factors participate in double-strand break repair during V(D)J recombination and immunoglobulin class-switch recombination. These alterations may therefore affect how DN cells respond to activation-associated genotoxic insults, disproportionately reinforcing the mutational processes detected in this compartment (**Fig. 2D**).

**Figure 3.**
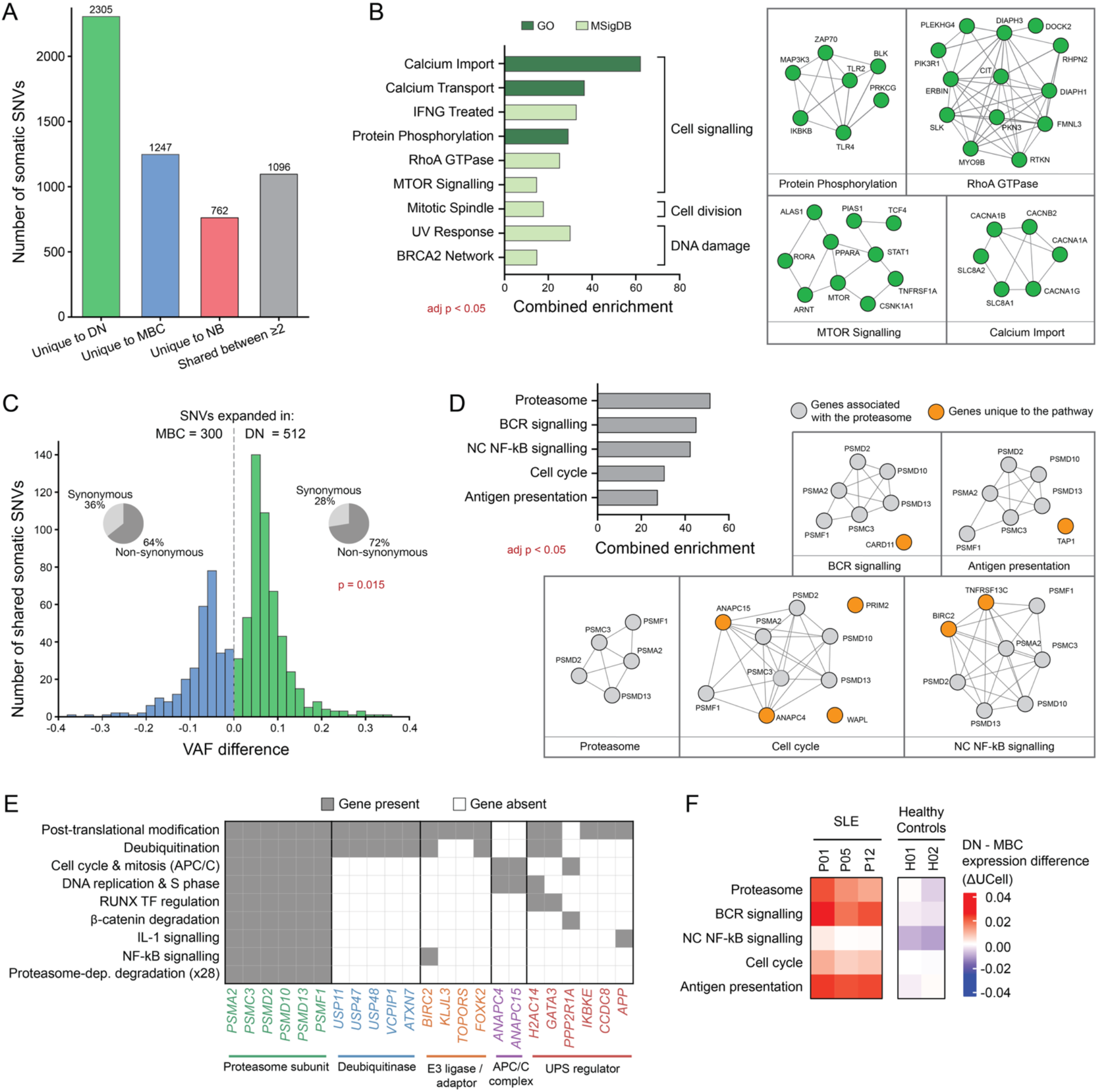
*De novo* and shared expanded variants contribute to the mutational burden and functional potential of DN B cells. **(A)** Number of somatic SNVs stratified by subset specificity. Shared SNVs are defined as variants detected at the same genomic coordinate, with the same base substitution, in at least two B-cell subsets from the same patient. **(B)** Pathway enrichment analysis of non-synonymous SNVs detected only in DN cells. Overrepresentation significance was assessed using a hypergeometric test, and pathways with FDR ≤ 0.05 were considered significantly enriched. Analyses were performed independently using GO Biological Process^40,41^, Reactome^42^, MSigDB^43^ C2, MSigDB C7 and MSigDB Hallmark. Combined scores reflect enrichment based on a weighted combination of odds ratio and *P* value^44^. Networks display representative genes from selected enriched terms and protein-protein interactions obtained from STRING^45^. **(C)** Distribution of VAF differences for SNVs shared between DN cells and MBC, calculated as DN VAF minus MBC VAF. Positive values indicate higher VAF in DN cells, consistent with relative expansion of variant-bearing cells in the DN compartment; negative values indicate higher VAF in MBC. The accompanying pie charts show the proportion of non-synonymous shared SNVs among variants with higher VAF in DN (green) or MBC (blue). *P* value was determined by chi-squared test. **(D)** Pathway enrichment analysis of non-synonymous shared SNVs with higher VAF in DN than in MBC. Pathways with FDR ≤ 0.05 were considered significantly enriched. Analysis was performed as in **(B)**. **(E)** Ubiquitin-proteasome system membership across enriched Reactome pathways for SNVs expanded in DN cells. **(F)** Heatmap of differential pathway activity between DN and MBC based on single-cell transcriptomic expression, assessed using UCell^46^ scores for gene sets corresponding to pathways enriched among DN-expanded shared SNVs.

We next asked whether variants shared across B-cell subsets showed evidence of preferential expansion in DN cells. Shared SNVs between NB and MBC or DN cells generally exhibited higher variant allele frequencies in the latter populations, and DN cells contained a significantly greater fraction of expanded shared mutations than MBC (p = 0.0002). Although these analyses do not resolve the precise developmental origins of DN cells, they are consistent with a model in which variants acquired before subset divergence subsequently undergo preferential expansion in the DN compartment. Direct comparison of SNVs shared between DN and MBC compartments showed that 63% had higher VAF in DN cells (**Fig. 3C**, main). Compared with MBC-expanded shared SNVs, DN-expanded SNVs contained a significantly higher proportion of non-synonymous substitutions (p = 0.015; **Fig. 3C**, insert), supporting enrichment of functionally consequential variants among clones expanded in the DN compartment.

Pathway enrichment analysis of DN-expanded SNVs identified proteasome function, B-cell receptor signalling, non-canonical NF-κB signalling, antigen presentation, and cell-cycle regulation as major affected processes (**Fig. 3D**). Altered genes within these pathways included various genes whose dysregulation is linked to B-cell autoimmunity (e.g., *CARD11*^11^, *TNFRSF13C/BAFFR*^37^, and *TAP1*^42^). Beyond these well-characterized genes, particularly notable was the recurrent involvement, across multiple patients, of genes linked to proteostasis. These not only included core proteasome subunits, but also several genes involved in ubiquitin conjugation, deubiquitination, and broader ubiquitin-proteasome system regulation (**Fig. 3D-E**). Given the central role of proteostasis in immune-cell activation and the ongoing therapeutic interest in this pathway in SLE^38,39^, these observations raise the possibility that somatic perturbation of proteostasis-related programs contributes to DN-cell persistence and behaviour.

A similar pattern was observed for indels. Indels shared between NB and either MBC or DN generally showed higher VAF in the differentiated subsets than in NB, consistent with preferential post-naïve expansion of indel-bearing clones.

The recurrent accumulation of somatic alterations in specific pathways across multiple patients suggested that variants converge on functional programs active in DN cells. To find phenotypic evidence that the pathways highlighted by the somatic variant analyses were indeed impacted in DN cells, we profiled PBMCs from three SLE patients in our cohort and two healthy reference controls by single-cell RNA sequencing. Across 35,176 B cells, SLE DN cells showed altered expression of the same pathways in the mutational analyses, including proteostasis, cell-cycle regulation, stress responses, and immune signalling (**Fig. 3F**). These findings provide orthogonal support for the relevance of pathway-level perturbations identified by somatic variant analyses in the DN compartment.

### DN cells harbour lymphoma-associated somatic mutations in SLE

Patients with SLE have a 2- to 7-fold increased risk of developing B-cell lymphomas relative to the general population^19^, raising the possibility that the disease and/or its management create conditions permissive for the acquisition or persistence of lymphoma-relevant lesions. We therefore asked whether DN cells in SLE harbour somatic mutations affecting genes recurrently altered in B-cell malignancies. Using a curated set of 144 high-confidence lymphoma-driver genes^47^, we identified 56 non-synonymous somatic SNVs and 4 indels affecting 45 “Tier 1” lymphoma-associated genes in DN cells (**Fig. 4A**). These mutations were detectable in the majority of patients (19/35 [54%]; **Fig. 4B**), with several individuals harbouring multiple lymphoma-associated mutations within the DN compartment, indicating that lymphoma-relevant genetic alterations are detectable within a restricted autoimmune B-cell pool in patients without overt lymphoma.

**Figure 4.**
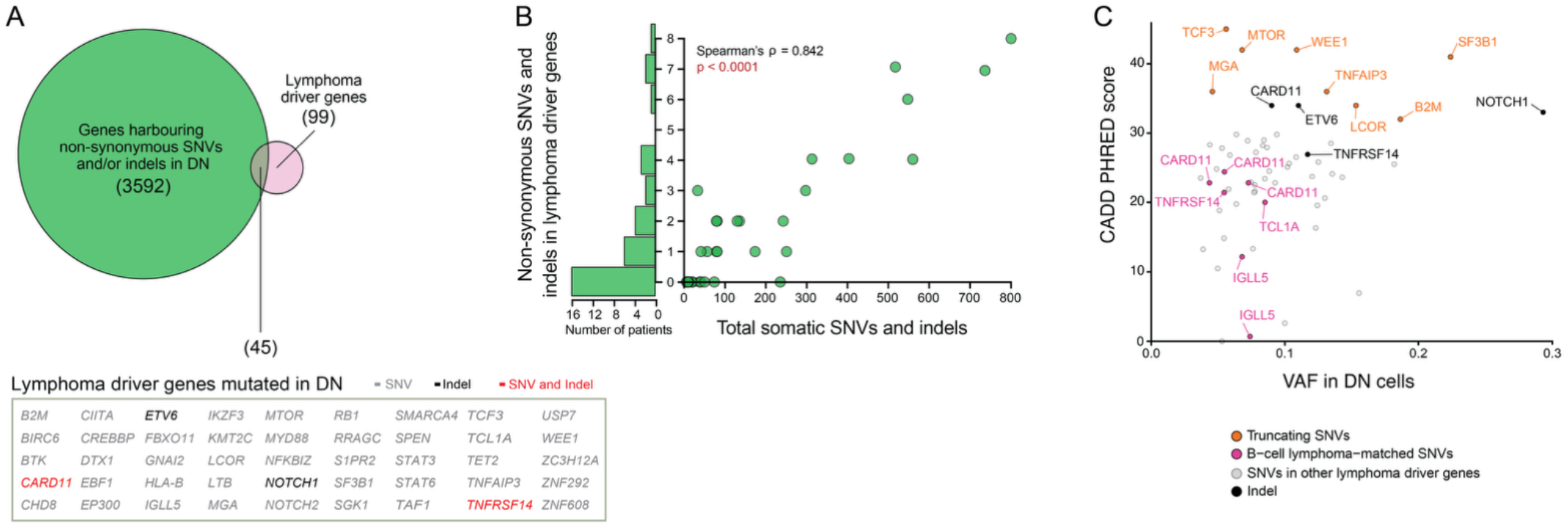
Lymphoma-associated somatic mutations are prevalent in DN B cells from patients with SLE. **(A)** Venn diagram showing the overlap between genes harbouring non-synonymous somatic mutations in DN cells, including SNVs and indels, and Tier 1 lymphoma-associated genes defined by Coyle et al.^47^. The inset lists the 45 lymphoma-associated genes mutated in DN cells across the cohort. **(B) *Left***, number of non-synonymous somatic mutations, including SNVs and indels, in lymphoma-associated genes detected in DN cells from each patient. ***Right***, association between the number of non-synonymous somatic mutations in lymphoma-associated genes and total somatic mutational burden in DN cells per patient. Associations were assessed using Spearman rank correlation; Spearman’s rho (ρ) and corresponding *P* value are shown. **(C)** Variant allele frequency (VAF) and predicted deleteriousness of DN mutations in lymphoma-associated genes. Scatter plot shows VAF versus CADD PHRED-scaled deleteriousness score^48^ for non-synonymous SNVs and indels in lymphoma-associated genes. Orange points denote truncating mutations annotated using the Ensembl Variant Effect Predictor (VEP)^49^; pink points denote mutations that exactly match, or are immediately adjacent to, amino-acid substitutions or mutations previously reported in B-cell lymphoma datasets from cBioPortal^50^; black dots denote indels. See **Methods** for details.

Both DN-specific and DN-expanded variants contributed to this lymphoma-associated mutational repertoire. The number of lymphoma-associated mutations in DN cells strongly correlated with total DN somatic mutational burden (ρ=0.842, p<0.0001; **Fig. 4B**), suggesting that lymphoma-associated mutations scale with the broader somatic diversification processes active in this compartment. VAFs of lymphoma-associated mutations were generally still low (median [Q1-Q3] of 0.078 [0.057 -0.112]), consistent with their presence in subclonal fractions of the DN compartment captured at the time of sampling (**Fig. 4C**).

Several mutations showed high predicted pathogenicity (CADD scores^48^; **Fig. 4C**), including truncating lesions in *MGA*, linked to enhanced MYC activity^51,52^, *TNFAIP3/A20*, linked to de-repressed NF-κB signalling^53^, and *B2M*, linked to immune evasion^54^. We also identified a *MYD88* mutation affecting the TIR domain, which contains the recurrent lymphoma-associated L265P hotspot. Indels showed a similar pattern, affecting established lymphoma-associated genes and further expanding the spectrum of potentially functional lymphoma-like lesions present in DN cells (**Fig. 4C**).

Together, these findings show that DN cells in SLE harbour somatic mutations affecting lymphoma-associated genes, including truncating lesions. These data connect chronic diversification of a pathogenic autoimmune B-cell compartment with mutational features associated with B-cell malignancy.

## DISCUSSION

Our study provides, to our knowledge, the first subset-resolved analysis of somatic mutational landscapes across circulating B-cell populations in systemic autoimmunity. By combining high-coverage exome sequencing of purified naïve, memory, and DN B cells, we identify DN B cells as the circulating B-cell compartment with the greatest degree of somatic genome diversification in SLE. DN cells carried the highest overall somatic mutational burden and showed evidence of ongoing mutation acquisition, preferential expansion of shared variant-bearing clones, and enrichment for lymphoma-associated alterations. Although prior studies have identified somatic variants in selected autoimmune B-cell contexts^11,12^, our subset-resolved design shows that somatic diversification is unevenly distributed across circulating B cells and is concentrated in the pathogenic DN compartment. We further integrate single-cell transcriptomic profiling to contextualize these mutations within disease-associated B-cell states and to support orthogonal validation of transcriptional programs associated with somatically diversified DN B cells. These findings extend the emerging view that somatic evolution can shape pathogenic immune states outside of cancer, particularly in chronic inflammatory disease settings^12,55,56^.

DN cells have been linked to defective tolerance, plasma-cell differentiation, autoantibody production, and disease activity^9,23,24^. Our data suggest that these cells also represent a genetically dynamic compartment that integrates the cumulative effects of chronic disease exposure. The lack of association with age argues against simple time-related accumulation as the main explanation for this pattern. Similarly, the absence of association with disease activity at sampling, cumulative disease activity, or flare frequency suggests that acute inflammatory burden alone does not account for the observed mutational landscape. Instead, the correlation with disease duration points to a longitudinal process in which repeated immune activation, proliferative history, oxidative stress, and treatment-related genotoxic exposures progressively shape the DN compartment. Mutational signature analysis supports this interpretation. DN cells carried signatures linked to oxidative damage, DNA repair-associated processes, and therapy-associated mutagenesis, including signatures associated with thiopurine exposure. Prior work in epithelial tumours showed that chronic azathioprine exposure can generate a distinct mutational signature whose contribution correlates with treatment duration^57^, providing precedent for the idea that immunomodulatory therapy can leave durable somatic footprints. Although our study examines non-malignant circulating immune cells rather than epithelial tumours, these observations support the broader concept that pathogenic B-cell populations can record mutational pressures encountered during chronic autoimmune disease. Because treatment history is closely intertwined with disease severity, duration, and clinical management, our analyses do not establish a causal role for any individual therapy. Nevertheless, the enrichment of treatment-associated signatures in DN cells, together with stratification by azathioprine exposure, suggests that immunomodulatory therapies may contribute to somatic diversification in circulating immune compartments. More generally, these data raise the possibility that chronic autoimmune disease creates a mutagenic environment in which inflammatory damage, replicative stress, and therapeutic pressure jointly increase the likelihood that pathogenic immune clones acquire and retain somatic alterations.

DN cells contained both subset-specific variants and shared variants that preferentially expanded within the DN compartment, indicating that this population is shaped by two complementary processes: ongoing mutation acquisition after entry into the DN state, and selective enrichment of pre-existing variant-bearing clones. The pathways affected by these variants converged on core features of B-cell function and pathogenicity implicated in SLE, spanning central activation cascades, alongside essential cellular maintenance programs, including antigen presentation, cell-cycle regulation, DNA damage responses, and proteostasis^58^. Although these pathways are well established in SLE biology, our data suggest that somatic alteration of key mediators within pathogenic B-cell compartments may provide an additional mechanism of disease dysregulation. In this way, somatic mutations may contribute to SLE heterogeneity beyond germline risk, transcriptional state, and environmental exposure. Particularly striking was the recurrent involvement of proteasome and ubiquitin-proteasome system-associated genes across multiple patients. Proteostasis has central roles in immune-cell activation, stress adaptation, and antigen processing, and this pathway is already being explored therapeutically in SLE^38,39^. Our findings raise the possibility that somatic perturbation of proteostasis-related programs contributes to DN-cell persistence or selective fitness in chronic inflammatory settings. Importantly, the altered genes extended beyond core proteasome components to include regulators of ubiquitin conjugation and deubiquitination, suggesting that upstream control of proteostasis may also be somatically disrupted in pathogenic B-cell states.

A major implication of our findings is the connection between systemic autoimmunity and lymphoma-associated somatic evolution. Patients with SLE have an increased risk of B-cell lymphoma^19,20^, and recent evidence suggests that immunosuppressive drug exposure may further contribute to lymphoma risk in autoimmune disease^59^. Yet the molecular basis of this association remains incompletely understood, and germline variants provide only a limited explanation^60^. Our data show that lymphoma-associated mutations are detectable across a substantial fraction of patients with SLE, affect genes recurrently altered in B-cell malignancies. Although these findings do not imply malignant transformation^61^, they suggest that pathogenic autoimmune B-cell populations can provide a substrate in which lymphoma-relevant lesions arise and persist before overt transformation^10,62^.

The strong correlation between lymphoma-associated mutations and total DN mutational burden further suggests that these alterations emerge as part of the same processes driving broader somatic diversification in DN cells. Their generally low VAFs are consistent with detection in subclonal fractions of circulating DN cells from individuals without clinically manifest lymphoma. However, low VAFs may also indicate that additional contextual requirements, such as antigenic stimulation, tissue localization, inflammatory niches, immune escape, or acquisition of cooperating lesions, are needed for variant-bearing DN cells to undergo more substantial clonal expansion. Together, these findings support a model in which chronic systemic autoimmunity and B-cell malignancy occupy overlapping evolutionary space. DN-like or age-associated/autoreactive B-cell programs have been implicated in lymphoid malignancies^10,62^, including extranodal lymphomas with transcriptional features reminiscent of aged/autoimmune B-cell states^10^. Our results add a genetic dimension to this connection by showing that autoreactive DN cells in SLE harbour mutations in lymphoma-associated genes. This suggests that chronic autoimmunity may create a permissive setting in which autoreactive B-cell states expand, accumulate somatic lesions and, in some cases, cross additional evolutionary thresholds toward malignant transformation.

Somatic diversification of DN cells may also contribute to SLE heterogeneity in the absence of lymphoma. DN cells may be particularly permissive for the phenotypic consequences of such mutations because they are activation-prone, exposed to inflammatory cues, and capable of differentiation toward antibody-secreting states. Somatic variants targeting core programs could fundamentally alter DN cell fate, disrupting how these cells respond to antigen and interact with T cells, while potentially re-shaping their survival under inflammatory stress, differentiation into plasma cells, and/or trafficking to tissues. In this way, somatic mutations may contribute not only to long-term lymphoma risk but also to differences in disease persistence, treatment response, organ involvement, or flare propensity. Because these variants can be detected from circulating B-cell populations, DN-focused profiling may eventually provide a minimally invasive strategy to monitor clonal features associated with high-risk or divergent disease trajectories.

Beyond the important insights highlighted above, this study establishes a foundational resource designed to move the field from broad discovery toward the targeted investigation of pathogenic immune evolution. By providing a high-depth map of purified B-cell subsets, we establish a high-confidence baseline of recurrently mutated genes and residues in non-malignant autoimmunity. This curated catalogue directly enables the design of targeted sequencing panels for longitudinal monitoring in clinical cohorts, offering a tool to determine if specific mutational signatures or clonal expansions can predict therapeutic response and disease flares. Furthermore, this resource provides a critical framework for the functional prioritization of somatic variants. By identifying alterations affecting residues recurrently mutated in malignancy and defining the transcriptional programs enriched in somatically diversified DN B cells, our data provide a roadmap for future studies modelling the impact of somatic mutations on B-cell fitness, signalling, and differentiation. Finally, this dataset serves as an essential comparative reference for both the immunology and oncology communities, providing the necessary context to determine whether these clonal trajectories represent SLE-specific pathology or generalized hallmarks of chronic immune activation and pre-malignant evolution.

In summary, we identify DN B cells as a compartment of enhanced somatic genome diversification in SLE and show that this pathogenic population harbours evolutionary features that overlap with those of lymphomas. Serving as a foundational resource for the study of systemic autoimmunity, our findings support a model in which chronic autoimmune disease is shaped by progressive somatic evolution within pathogenic immune compartments. This framework opens a new perspective on disease heterogeneity and long-term complications, providing a critical lens through which to investigate the biological continuum between autoimmunity and malignancy.

## METHODS

### Study participants

Adult patients included in this study provided written informed consent in accordance with institutional review board protocol 22-5096, approved by the University Health Network Research Ethics Board.

### Sample collection

Whole blood and buccal swabs were collected at study entry by clinical staff at the Toronto Lupus Clinic (Toronto Western Hospital, University Health Network). Peripheral blood mononuclear cells (PBMCs) were isolated within 4h of blood draw using Ficoll-Paque^TM^ Plus (Sigma-Aldrich, St. Louis, United States) density-gradient centrifugation in Sepmate^TM^ Tubes (Stemcell Technologies, Vancouver, Canada), according to the manufacturer’s instructions. PBMCs were frozen at -80°C for 24 h in Bambanker^TM^ Serum-Free Freezing Media (GC Lymphotec Inc., Tokyo, Japan) and subsequently transferred to liquid nitrogen for long-term storage. Frozen PBMC from healthy controls were purchased from Stemcell Technologies and stored in liquid nitrogen. Buccal swabs were collected in OraCollect tubes (Cat# OCR-100, DNA Genotek, Ottawa, Canada) and stored at 4^°^C for a maximum of one month prior to DNA extraction.

### Cell sorting

Live-cell sorting was performed to simultaneously isolate naïve B-cells (NB; CD19^+^IgD^+^), memory B-cells (MBC; CD19^+^IgD^−^CD27^+^), and double-negative B-cells (DN; CD19^+^IgD^−^CD27^−^), as well as monocytes (CD19^−^CD14^+^) and hematopoietic stem cells (HSC; CD19^−^CD34^+^). To this end, cryopreserved PBMCs were rapidly thawed and washed once in DPBS containing 40% FBS. Cells were stained with fluorescently conjugated monoclonal antibodies and SYTOX™ Blue for viability. Cells were washed in DPBS + 5% FBS, filtered through a 70 µm cell strainer and sorted immediately on a FACSAria^TM^ Fusion Cell Sorter (BD Biosciences, Franklin Lakes, USA). Sorted cells were frozen at -80^°^C in Bambanker^TM^ Serum-Free Freezing Media (GC Lymphotec Inc.) and stored until DNA extraction.

### DNA extraction and whole-genome amplification

DNA was extracted from buccal swabs and sorted cell samples containing ≥2,000 cells (resuspended in PBS) using the QIAamp DNA Mini Kit (Cat# 51304, Qiagen, Hilden, Germany), following the manufacturer’s protocol for blood and body fluids. For buccal samples, an additional pre-incubation step was included: tubes were incubated for 2 h at 50°C in a heat block prior to proceeding with the QIAamp protocol. For sorted cell samples with <2,000 cells, DNA was amplified directly from cell lysates using the REPLI-g Single Cell Whole Genome Amplification Kit (Cat # 150343, Qiagen, Hilden, Germany), according to the manufacturer’s “Protocol 1” recommendations. Purified Genomic DNA from sorted B-cell samples with ≥2,000 cells were amplified according to the manufacturer’s “Protocol 2” recommendations. Samples containing fewer than 260 sorted cells (n=4 NB samples) were excluded from sequencing and downstream analysis.

### Whole-exome sequencing

Whole-exome sequencing (WES) was performed at the Ontario Institute for Cancer Research (Toronto, Canada). A total of 206 samples were subjected to WES. These included NB, MBC, and DN B cells from each of the 35 patients with sufficient sorted cell counts for a given cell subset, comprising 31 NB samples and 35 MBC and DN B samples. Matched HSCs and monocytes were sequenced as non-B-cell lineage controls (n=35, respectively), and matched buccal samples were sequenced as patient-specific matched normals for somatic variant calling (n=35). Libraries were prepared using the xGen™ Exome Hyb Panel v2 (Integrated DNA Technologies, Coralville, USA) and sequenced on an Illumina NovaSeq X Plus system (Illumina, San Diego, USA) to a target coverage of 200x.

### Read alignment

Raw reads were processed with the nf-core pipeline Sarek^63^ (v3.2.3). Reads were aligned to the GRCh38 reference genome (GATK Resource Bundle) using BWA-MEM2^64^, with alignment, duplicate marking, and base quality score recalibration performed using Sarek default parameters. Analyses were restricted to regions targeted by the xGen™ Exome Hyb Panel v2.

### SNV calling and filtering

Germline variants were identified using GATK HaplotypeCaller^65^ to support germline filtering and assessment of cross-sample contamination. Somatic SNVs were detected using three callers: Mutect2^66^, Strelka2^67^, and CaVEMan^68^, each run in paired mode using the corresponding buccal swab sample as the matched normal. Mutect2 and Strelka2 were run as part of the Sarek pipeline (v3.2.3) using default settings. CaVEMan (v1.15.5) was run independently, and variant calls were processed with cgpCaVEManPostProcessing (v3.1.6), which annotates variants as PASS/FAIL based on multiple technical criteria.

To reduce false positive variants, several filtering steps were performed. First, only variants labelled as “PASS” by each caller were retained. Second, variants detected by ≥2 of the 3 callers were kept for downstream analysis and were subjected to further filtering based on an allele depth (AD) threshold of ≥5 and a total depth (DP) threshold of ≥20. Additional filters were applied according to previously published best practices^69^ to remove variants with evidence of pronounced strand bias, homopolymer-associated artifacts, and localization in low-mappability or low-complexity regions. Outlier samples were identified using Mahalanobis distance computed from the total somatic SNV burden, extracted DNA mass, and sorted cell count. Outlier samples (P16_NB, P13_CD34, P20_CD34, P21_CD34) were removed prior to downstream analysis.

Variant annotation was performed using ANNOVAR^70^ (with Gencode v45) and VEP^49^ (Ensembl release 113). SNVs labelled as “synonymous SNV” in the ExonicFunc.ensGene annotation field were classified as synonymous; all other coding variants were considered non-synonymous. Canonical transcript assignments and predicted consequences were taken from VEP annotations.

### Indel calling and filtering

Germline indels were identified using GATK HaplotypeCaller to support germline filtering and assessment of cross-sample contamination. Somatic indels were detected using three callers: Mutect2, Strelka2, and Pindel^71^, each run in paired mode using the corresponding buccal swab sample as the matched normal. Mutect2 and Strelka2 were run as part of the Sarek pipeline (v3.2.3) using default settings. Pindel (v3.2.0) was run using the cgpwxs pipeline (v3.1.6; https://github.com/cancerit/dockstore-cgpwxs), which annotates indels as PASS/FAIL based on multiple technical criteria.

To reduce false positives, only indels labelled as PASS by each caller were retained. Indels detected by at least two of the three callers were then kept for downstream analysis and further filtered using an allele depth threshold of ≥5 and a total depth threshold of ≥20. Additional filters were applied to remove indels located in UCSC Unusual Regions and RepeatMasker regions downloaded from the UCSC Genome Browser^72^. Outlier samples were identified using Mahalanobis distance computed from the total somatic indel burden, extracted DNA mass, and sorted cell count. Outlier samples (P16_NB, P13_CD34, P20_CD34, P21_CD34, P24_Memory) were removed prior to downstream analysis.

Indels were annotated using VEP, and canonical transcript assignments and predicted consequences were taken from VEP annotations.

### Mutational signature analysis

Mutational signature analysis was performed using the MutationalPatterns R package (v3.10)^73^. Because mutational signature deconvolution becomes unreliable when mutation counts are low, we restricted analysis to samples harbouring more than 50 high-confidence somatic SNVs. For each eligible sample, COSMIC single-base substitution (SBS) signatures included within the MutationalPatterns package were fitted using the function *fit_to_signatures_bootstrapped(method = “strict”)*, which applies iterative bootstrapping to improve robustness of signature assignment. Comparative analyses of mutational processes were limited to DN and MBC subsets, as only three NB samples exceeded the 50-mutation threshold required for inclusion.

### Identification of unique and shared mutations

To determine with high confidence whether somatic alterations were unique to, or shared among, B-cell subsets within each patient, we used a two-step approach, applying the same two-step approach independently on SNVs and indels. In the first step, we defined a set of candidate B-cell mutations: variants were initially compiled from all three B-cell subsets (DN, MBC, and NB) if they were detected by at least two of the three somatic callers after outlier removal and all upstream filtering. To ensure that only mutations arising within the B-cell lineage were carried forward, we excluded any variants detected in matched HSC (CD34^+^) and monocyte (CD14^+^) populations. These populations were sequenced in a patient-specific manner as non-B-cell lineage controls, enabling the removal of mutations originating in hematopoietic progenitors or myeloid cells.

Second, after establishing the B-cell-specific candidate set, we performed high-resolution presence/absence calling for each mutation across DN, MBC, and NB subsets by directly inspecting raw sequencing reads. For every site at which an SNV or indel was identified in at least one B-cell subset from a given patient, we examined per-base pileups using perbase (https://github.com/sstadick/perbase). To ensure comparability across subsets, each locus was required to have at least 50× perbase-reported coverage in all three B-cell populations. A mutation was considered present in a given subset if the perbase-derived variant allele frequency (VAF) was ≥0.005, equivalent to 1/200. This conservative threshold was selected to enable detection of low-frequency somatic variants while limiting spurious presence calls.

Analyses of subset-unique mutations were performed in the 30 SLE patients with non-outlier variant-calling results for all three B-cell subsets. Five patients were excluded: four had insufficient NB cell numbers for WES, and one displayed NB outlier status based on Mahalanobis distance. For these 30 patients, mutations meeting the perbase presence criterion in exactly one subset and absent from the remaining two subsets, defined as VAF <0.005, were classified as unique.

Shared mutations were identified using the same perbase coverage and VAF thresholds but requiring a mutation to be present in both subsets being compared. DN–MBC shared mutations were assessed in all 35 patients for whom both compartments were sequenced, whereas DN–NB shared mutations were evaluated only in the 30 patients with high-quality NB data. Mutations satisfying the perbase coverage and VAF criteria in both subsets were designated as shared.

### Overrepresentation analysis

Overrepresentation analysis (ORA) was performed using GSEAPY^74^ (v1.1.8) with pathway annotations from the following databases: GO Biological Process^40,41^ (2025 version, Human), Reactome^42^ (v89), MSigDB^43^ C2 curated gene sets (C2.CPG & C2.CP), MSigDB C7, and MSigDB Hallmark. For MSigDB C2, C7, and Hallmark gene sets, GMT files from database version 2024.1.Hs were used. The background gene set comprised the 19,766 genes targeted by the xGen™ Exome Hyb Panel v2, ensuring that enrichment statistics reflected the capture design of the sequencing assay. Pathways were considered significantly enriched if they met an adjusted *P value* threshold of ≤0.05. All analyses were conducted using default ORA settings in GSEAPY. Where applicable, Reactome’s hierarchical structure was used to assign pathway categories. Network diagrams were created by taking representative genes selected from enriched terms, with protein–protein interactions obtained from the STRING database^45^.

### Single-nuclei RNA sequencing

Cryopreserved PBMCs were thawed, stained, and sorted as described above. Sorted cells were pooled to enrich for MBC and DN, where ∼8×10^4^ MBC and DN cells (CD19^+^, IgD^-^) were combined with ∼1×10^4^ NB cells (CD19^+^, IgD^+^) and ∼1×10^4^ CD19^-^ cells. A total of approximately 1×10^5^ live cells were processed at the Princess Margaret Genomics Centre using the Chromium Next GEM Single Cell Multiome ATAC + Gene Expression (10X Genomics, Pleasanton, USA) according to the manufacturer’s CG000365 Demonstrated Protocol for low-cell-input nuclei isolation. Gene Expression libraries were sequenced separately on Illumina’s NovaSeq X with the following run parameters: read 1 – 28 cycles, read 2 – 90 cycles, index 1 – 10 cycles and index 2 – 10 cycles.

### Single-nuclei RNA-sequencing data processing

Raw single-nuclei RNA-sequencing data were processed using the Cell Ranger Arc^75,76^ (v2.0.2) pipeline, aligning to the bundled GRCh38 (v2020-A-2.0.0) genome to produce feature-barcode matrices for downstream analysis with *Seurat*^77^ (v4.4.0) in *R* (v4.3.1). Quality control included filtering cells with unique molecular identifier (UMI) counts more than three median absolute deviations from the median, gene counts above three median absolute deviations from the median or below 200 genes, and mitochondrial read percentage greater than 20%. Cell cycle scoring was carried out using the bundled `*cc*.*genes*.*updated*.*2019*` gene sets, and *SCTransform* (*vst*.*flavor=“v2”*) was then used to regress out effects of cell cycle, mitochondrial percentage, number of detected genes, as well as number of UMIs per cell.

Cells were clustered on the shared neighbourhood graph using the first 30 principal components across resolutions ranging from 0 to 1 in increments of 0.1. Then, Seurat label transfer using *Triana et al. (2021)*^78^ reference dataset, together with *Azimuth*^79^ annotation using the bundled PBMC reference dataset, provided two orthogonal sources of initial cell-type annotation. These annotations were combined with cluster marker genes identified using Seurat’s ‘*FindAllMarkers’* function to select the optimal clustering resolution for isolating B-cell-containing clusters. Once B-cell clusters were identified for each sample, a second round of cell typing was performed using similar steps, together with visualization of key marker genes, including *ITGAX, ZEB2, TRAF5*, and *ZEB1*, using *clustree*^80^ (v0.5.1) to determine the optimal clustering resolution for DN-cell annotation.

All samples were merged, and *Harmony*^81^ (v1.2.3) was used to integrate samples and generate a UMAP visualization using 30 components. The *FindConservedMarkers* function of Seurat was used to identify markers conserved across samples for each cell type, and the top 10 unique markers were visualized. UCell^46^ (v2.6.2) was used to score gene sets derived from msigdbr^82^ (v25.1.1) at a per-cell level. Median scores were calculated per sample and per cell type for G1-phase cells, and differences between DN cells and MBCs were visualized as heatmaps using *ComplexHeatmap*^83^ (v2.18.0).

### Statistical methods

No statistical methods were used to predetermine sample size. The study included all available consented SLE patients with sufficient sorted cell numbers and sequencing data passing quality-control criteria. Experiments were not randomized, and investigators were not blinded to sample identity during experimental processing or analysis. Unless otherwise stated, statistical analyses were performed at the patient or patient-derived subset level, as indicated in each figure legend. For comparisons across B-cell subsets, analyses were restricted to patients with matched evaluable NB, MBC and DN samples unless otherwise stated. Each patient contributed one value per B-cell subset, and no technical replicates were treated as independent biological observations. Associations between continuous variables were assessed using two-sided Spearman rank correlation. Comparisons of somatic mutational burden across B-cell subsets were performed using non-parametric tests followed by pairwise Dunn’s tests for multiple comparisons. Differences in categorical proportions were assessed using chi-squared tests or Fisher’s exact tests, as appropriate. Pathway overrepresentation analyses were performed using hypergeometric tests, with multiple-testing correction by the Benjamini–Hochberg false-discovery rate. Pathways with adjusted *P* ≤ 0.05 were considered significant. Unless otherwise stated, all statistical tests were two-sided, and exact *P* values are reported where appropriate. Data are summarized as median and interquartile range, or as otherwise indicated in the figure legends.

## CODE AVAILABILITY

All custom scripts and workflows used for preprocessing and variant calling are available at https://github.com/GaitiLab/sle-bcell-mutation-landscape.

## DATA AVAILABILITY

De-identified human patient FASTQ files are being deposited at the European Genome–phenome Archive (EGA).

## ACKNOWLEDGEMENTS

We gratefully acknowledge the patients who generously donated their time and samples for this study, as well as the dedication of the clinic staff involved in patient recruitment and sample collection. We thank the core facilities at the Ontario Institute for Cancer Research (OICR), Princess Margaret Genomics Centre, and the SickKids–UHN Flow Cytometry Facility for their technical support. Z.T. is supported by the Department of Medicine, University of Toronto, and the Murray B. Urowitz Chair in Lupus Research. L.V. is supported by a Michael Smith Health Research Foundation Scholarship (SCH-2022-2607), the Canada Foundation for Innovation John R. Evans Leaders Fund (43630) and Innovation Fund (46036), and a Canadian Institutes of Health Research Project Grant (180613). F.G. is supported by the Princess Margaret Cancer Foundation, an Ontario Institute for Cancer Research Investigator Award (IA-1-025), the Natural Sciences and Engineering Research Council of Canada (RGPIN-2023-05535), the Canadian Cancer Society (708077), the Lupus Research Alliance (1077421), and the Cancer Research Society (1050446).

## AUTHORSHIP CONTRIBUTIONS

F.G., L.V., and Z.T. conceived and designed the study. D.B., L.P.W-G., J.E.W., D.D.G., and Z.T. recruited patients to the study and collected clinical data. L.C., X.L., and J.I.N. performed experiments. Y.S., P.K., L.C., L.P., X.L., A.A., L.V., and F.G. analyzed the data. Y.S., P.K., L.C., L.P., X.L., L.V., and F.G. interpreted the analysis results. Y.S., P.K., L.C., L.P., L.V., and F.G. wrote the manuscript with comments and contributions from all authors. All authors reviewed and approved the final manuscript.

